# Multi-species benchmark analysis for LC-MS/MS validation and performance evaluation in bottom-up proteomics

**DOI:** 10.1101/2023.08.28.555075

**Authors:** Tobias Jumel, Andrej Shevchenko

## Abstract

We present an instrument-independent benchmarking procedure and software (LFQ_bout) for validation and comparative evaluation of the performance of LC-MS/MS and data processing workflows in bottom-up proteomics. It enables back-to-back comparison of common and emerging workflows, e.g. diaPASEF or ScanningSWATH, and evaluates the impact of arbitrary, inadequately documented settings or black-box data processing algorithms. The procedure enhances the overall performance and quantitative accuracy while enabling the detection of major error types.

## INTRODUCTION

Data-independent acquisition (DIA)^1^ is becoming increasingly popular in bottom-up proteomics, while more dedicated instrumentation and software are becoming available (reviewed in ^2–4^). The performance and quantitative accuracy of DIA workflows is typically benchmarked using multiple mixtures of total proteome digests from 2-4 species^5^. Analyzing the series of samples with pre-defined fold–changes between individual proteomes allows comprehensive assessment of the quantification accuracy at the proteome-wide scale ^5,6,15,7–14^.

However, we have noticed that the output of benchmarking experiments is typically presented as protein group-level scatter or density graphs together with a single summary statistic related to the difference between expected and measured fold-changes. Within this framework, which is lacking an appropriate set of summary statistics and thresholds, it is difficult to detect errors and to perform data-based accuracy evaluations and performance comparisons.

To streamline workflow evaluation and optimization, we combined multi-species measurements with an R script providing the required visualizations and summary statistics. Part of the procedure is differential expression analysis of protein groups similar to previously described protocols ^12,16^. Systematic application of our benchmark workflow to own and external datasets acquired using common proteomics platforms identified five major sources of quantification errors.

We demonstrate that the impact of this straightforward, yet comprehensive benchmarking protocol extends beyond routine quality control and is critical for data integrity, inter-laboratory consistency and interpretation transparency of proteomics experiments.

## EXPERIMENTAL PROCEDURES

### Data and Code Availability

The benchmark analysis script and supporting materials are available at https://github.com/t-jumel/LFQb. Raw data are available at MassIVE-KB (https://massive.ucsd.edu/ProteoSAFe/static/massive.jsp) with the identifiers MSV000090837 and MSV000090832 (Supplementary Figures S1 and S2). External raw data reanalyzed in this work (not from QE-HF instrument) were from PXD028735 (https://proteomecentral.proteomexchange.org/cgi/GetDataset?ID=PXD028735).

### Benchmark Samples and Experiment Design

Multi-species sample mixtures ^5^ with expected log2 fold changes (A/B) of 0 for human, +1 for yeast, and - 2 for *E.coli* were prepared using Pierce HeLa Protein Digest Standard (Thermo Fisher Scientific), MS Compatible Yeast Protein Extract, Digest (Promega GmbH, Walldorf, Germany), and MassPREP E. coli Digest Standard (Waters Corporation, Milford, USA). Sample A consisted of 45 ng *E. coli*, 270 ng yeast, and 585 ng human protein digest while sample B consisted of 180 ng *E. coli*, 135 ng yeast, and 585 ng human protein digest per injection, calculated from the total source digest amount indicated by the manufacturers. Both samples were injected with a total load of 0.9 µg on column at a concentration of 0.18 µg/µl in 0.2 % formic acid and analyzed in triplicates in a block-randomized fashion ^17^. The normalisation benchmark data were acquired using 1 µg sample A on column and 0.7 µg sample B on column, in 2 technical replicates, using 30-minute elution gradients, and both DDA and DIA acquisitions in a block-randomized fashion. The diaPASEF and other non-QE-HF raw data were from PXD028735 ^14^.

### LC-MS

The LC-MS setup (all from Thermo Fisher Scientific, except where indicated) consisted of an UltiMate 3000 UHPLC system equipped with an Acclaim PepMap precolumn (100 µm x 20 mm, C18, 5 µm, 100 Å) and a 50 cm µPAC pillar array column. 5 µl samples were injected at a flow rate of 5 µl/min and eluted using a 2-sloped linear 90 min gradient (all steps 0.5 µl/min, 0.1 % FA) from 0 - 17.5 % ACN in 60 min (two thirds of the gradient length) and 17.5 - 35 % ACN in 30 min (one third of the gradient length). The LC was coupled to a QE-HF via a μPAC Flex iON interface plus nESI emitter (20 µm, 5 cm, Fossiliontech, Madrid, Spain). Electrospray voltage was 2.5 kV, transfer capillary temperature was 280 °C, and S-lens RF level was 50 %.

### Proteomics Data Acquisition and DDA Raw Data Processing

Data-independent acquisition (DIA) consisted of a full MS scan (m/z range of 395-955, R*_m/z_* _200_ 30000, 3 x 10^6^ automatic gain control, 55 ms injection time, centroid mode) and a series of 31 MS2 scans (R*_m/z_* _200_ 30000 m/z, 1 x 10^6^ automatic gain control, 55 ms injection time, centroid mode, 18 m/z isolation window width, normalized collision energy 24, fixed first mass 100 m/z) covering 400 – 950 precursor m/z and 100 – 2000 fragment m/z after demultiplexing with staggered DIA windows ^18^. Raw files were demultiplexed and converted to mzML using MSConvert v3.0.2 ^19^ and processed with DIA-NN v1.8, using predicted spectral libraries ^6^. Visualization of DIA data was performed in Skyline-daily v21.2.1.514 ^20^.

Data-dependent acquisition (DDA) included an MS1 scan (m/z range of 350-1700, R*_m/z_* _200_ 60000, 3 x 10^6^ automatic gain control, 55 ms injection time, profile mode) followed by Top15 data-dependent MS2 (R*_m/z_* _200_ 15000, 1 x 10^5^ automatic gain control, 50 ms injection time, centroid mode, 1.6 m/z isolation window width and offset of +0.2 m/z, normalized collision energy 24, fixed first mass 100 m/z, dynamic exclusion 20 s, all charges excluded except 2-5). DDA data were analyzed with MaxQuant v1.6.17.0 ^21^ and MSFragger v3.1.1 / FragPipe version 14.0 ^22^ with default settings without MBR.

### DIA Raw Data Processing

The “default” settings of DIA-NN v1.8 have been adjusted to ensure that the MBR is disabled and that the appropriate precursor and fragment m/z ranges for QE-HF are 400-950 and 100-2000 respectively. “Optimized” DIA-NN settings include: --cut K*,R* --var-mods 1 --var-mod UniMod:35,15.994915,M --double-search --individual-mass-acc --individual-windows --smart-profiling --pg-level 2 --species-genes --peak-center --no-ifs-removal --no-quant-files --report-lib-info --il-eq --matrix-qvalue 0.005 --nn-single-seq. For the search, we used canonical SWISS-PROT proteome databases derived from UniProt as of 20.08.2021 for the 3 species within the benchmarks (human - UP000005640, yeast - UP000002311, and E.coli - UP000000625) as well as the MaxQuant contaminant database ^21^. Raw data from PXD028735 ^14^ were analyzed with optimized DIA-NN analysis settings as listed above and with precursor and fragment m/z ranges of 399-1201 and 50-2000 for diaPASEF and TTOF5600, 399-901 and 50-2000 for QE-HF-X, Scanning SWATH, as well as 399-1201 and 100-1500 for TTOF6600Swath.

#### Benchmark Analysis Script

The DIA-NN-style protein group and precursor matrices were analyzed with our in-house developed R script available at https://github.com/t-jumel/LFQb. The script reported an average and median CV, asymmetry factors, confusion matrix summary statistics, and statistics related to the log2 fold change values. Briefly, entries matching MaxQuant contaminants or multiple species were removed. Entries were considered as identified, if data completeness exceeded 50 %, i.e. a value was reported in at least 2 out of 3 replicates in both Sample A and B. Identified entries, further characterized by a coefficient of variation (CV) below 20 % in both Sample A and B, were considered as quantified.

The asymmetry factor ^23^ was derived from the density function of log2 fold-change values by dividing the left and right distances between the center line and the x-values at 10 % of the maximum height. An asymmetry factor <1 indicated underestimation of fold changes and a value >1 indicated overestimation. Thresholds for reaching an undesirable degree were 0.5 and 2, respectively.

The quantified protein groups were further subjected to differential expression analysis using Limma v3.50.0 based on log2-transformed intensities and robust empirical Bayes statistics. Protein groups with a BH-adjusted p-value less than 0.01 and a log2 fold-change greater than 0.5 were considered as up- or down-regulated. By comparing these measurement results with expectations based on the sample mixtures, we obtained confusion matrix summary statistics ^24^. The approach was similar to that used in ^7^ and ^12^. Our focus was on the number of true positives (TP) and the false discovery rate, here called deFDR (differential expression FDR). The deFDR was calculated according to the confusion matrix by FP / (FP+TP). Other confusion matrix statistics were the sensitivity or true positive rate calculated by TP / (FP+TP), and the specificity or true negative rate calculated by TN / (TN+FP).

Secondary summary statistics, which acted as additional indicators, can be explained as following: The “Accuracy” summary statistic referred to the average distance between expected and measured log2 fold changes and was not associated with confusion matrix counts. The “Dispersion” was the average distance between measured log2 fold changes and their respective medians. The “Trueness” was the sum of the distances between measured median and expected log2 fold-changes for the species involved.

## RESULTS

### Limitations of Current Benchmark Procedures

Quantification accuracy in untargeted, label-free, bottom-up proteomics is not rigorously defined. While typical multi-species benchmarks support the evaluation of quantitative accuracy, we argue that common reporting formats are insufficiently informative (Figures 1A and 1B). In particular, the use of a single scatterplot and a single summary statistic of the distance between measured and expected log2 fold changes does not reflect the presence of multiple different sources of error. Based on our experience and extensive benchmarks, including re-processed raw data from repositories ^14^, we have identified five major sources of error in bottom-up proteomics (Figure 1C). These are:

**Figure 1:**
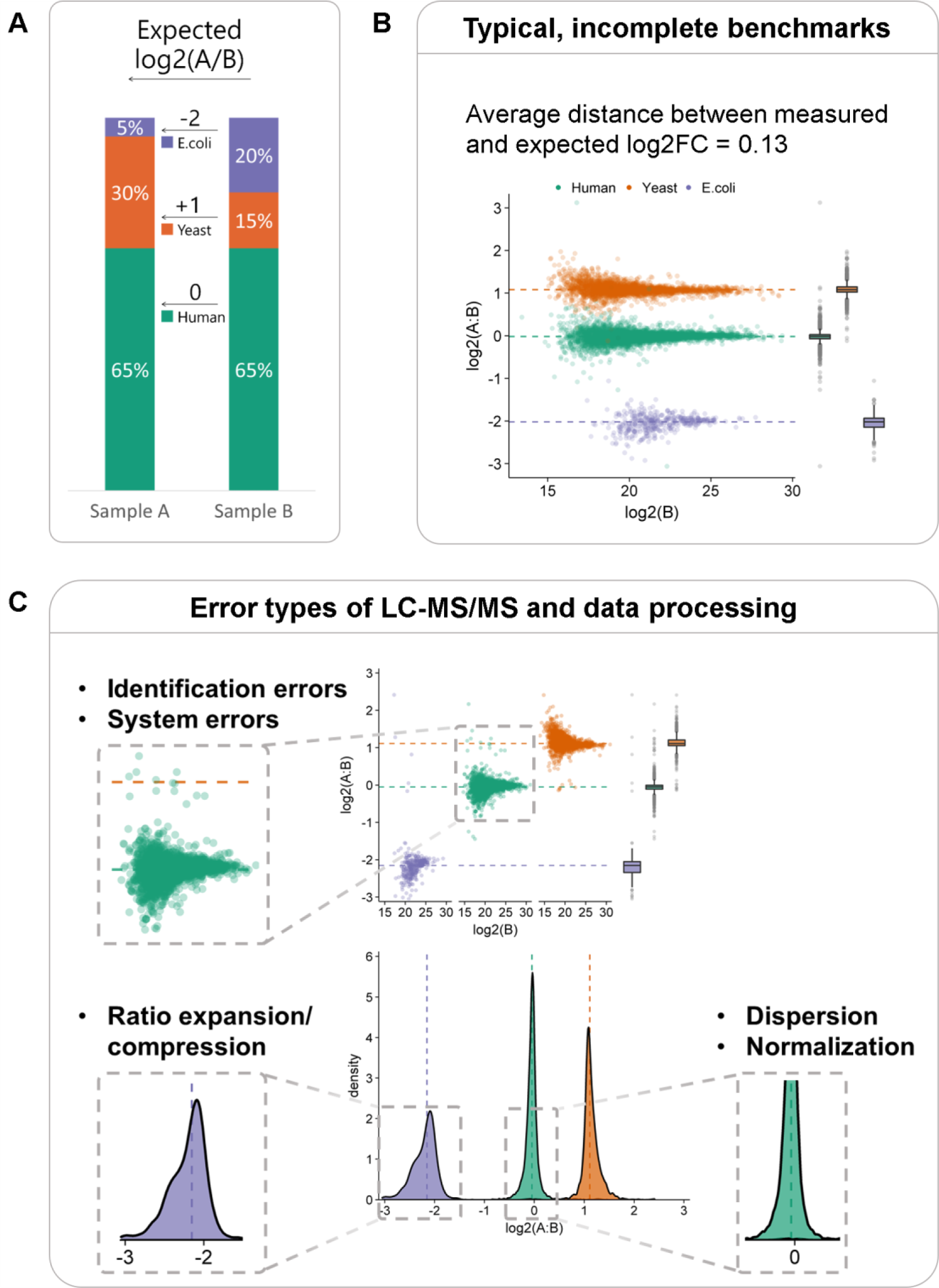
Limitations of current bottom-up proteomics benchmark protocols. **A)** Multi-species sample types used for QE-HF measurements and data from PXD0287352. **B)** Benchmark results typically consist of scatter plots and differences between expected and measured fold change values of protein groups, providing limited insight into data quality. **C)** Typical types of errors that benchmarks should detect, exemplified by QE-HF DIA data analysed with default settings in DIA-NN v1.8. While facet and density plots illustrate most error types, their detection should primarily be performed using summary statistics.

I) incorrect identifications resulting from mismatched peptide sequences in the database; ii) quantitative dispersion, typically expressed as coefficient of variation (CV). Low dispersion (high precision) is a prerequisite for high accuracy; iii) non-systematic over- or underestimation of fold changes between conditions (ratio expansion and ratio compression, summarized in this paper as “ distortion”); iv) MS system errors leading to erroneous peak intensities in the raw data, possibly related to the instrument control software (Supplementary Figure S4); v) cross-run normalisation errors during data processing. (Supplementary Figures S1 and S2).

We reasoned that multi-species benchmarks could be used to ensure the absence of system and normalisation errors, and to recognize and reduce errors related to dispersion, distortion and identification (Figure 2B-2). Error evaluation (Figure 2), can be performed using our R script and the benchmarked workflow is classified as either accurate or inaccurate. In parallel, we evaluated and compared workflow performances using the number of true positives or precise protein group quantifications instead of ID numbers.

**Figure 2:**
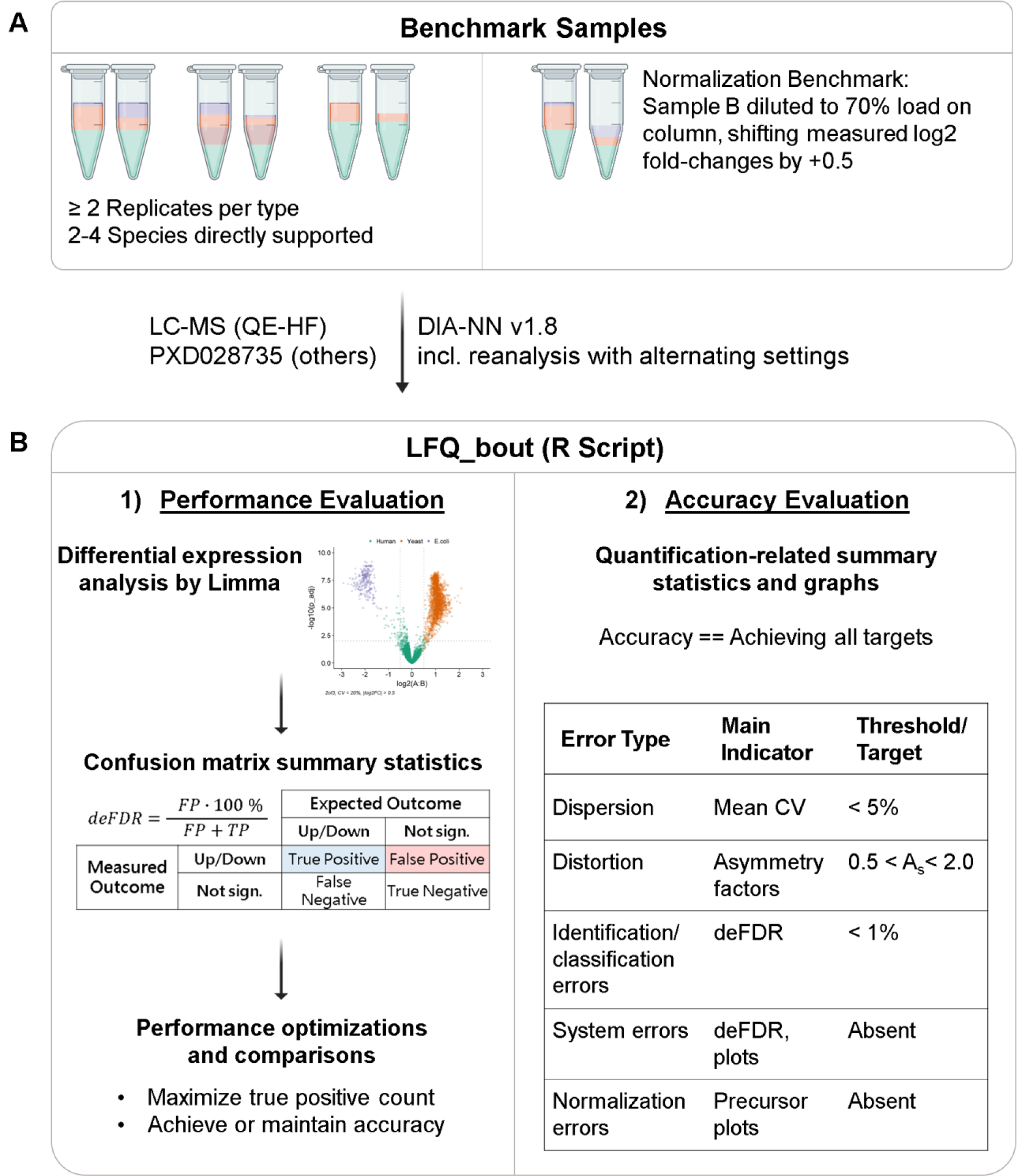
Multi-faceted benchmarks for complete validation and evaluation of untargeted bottom-up proteomics. **A)** The R script can process data from all common benchmark samples. A new variant requires the measurement of a diluted sample to validate the normalisation algorithms. **B1)** Quantified protein groups are processed with differential expression analysis to provide essential summary statistics to evaluate performance. **B2)** Overview of minimum summary statistics to cover all error types and suggested thresholds for the particular benchmark type. Only workflows that pass all criteria are considered “accurate” in parallel with performance evaluation.

A core component of the benchmark workflow is the differential expression analysis of protein groups. Based on the measured fold changes and adjusted p-values, protein groups are classified as upregulated, stable or downregulated and compared with the expectation defined by the sample mixtures. In this way, each protein group quantification is further classified as true (or false) positive (TP / FP) or true (or false) negative (TN / FN).

The number of protein groups correctly measured as up-regulated (TP count) is used as the main workflow performance indicator in parallel with the accuracy evaluation. The TP count is always accompanied by the classification error rate (deFDR = FP/(FP+TP)). The deFDR allows the user to detect whether workflows overestimated the number of identified proteins because of a poor identification stringency or might lead to increased errors in differential expression analysis due to ratio expansion or system errors. By using a 1 % deFDR threshold, these workflows can be classified as inaccurate.

Other summary statistics describing dispersion and distortion (Figures 1 and 2) are derived directly from the measured log2 fold changes of the protein groups. The graphical output produced for both protein groups and precursors also detects unexpected system and normalisation errors that are only visible at the precursor level.

We proposed that benchmark results are best evaluated using true positive counts for comparative rankings, but only TP counts with associated deFDR values below the threshold are considered meaningful. Workflow optimization should aim to maximize the TP count while meeting all accuracy requirements as shown in Figure 2B-2.

### Benchmark-Guided Workflow Evaluation and Optimization

We exemplify the value of our benchmark procedure by comparing QE-HF data processed 4 times under different DIA-NN settings. The four quantification strategies we tested used different settings that are supposed to control how chromatographic peak boundaries are recognized and peaks integrated, but the true impact on the performance and accuracy of the workflow was unknown. The four quantification strategies are combinations of either the “high precision” or “high accuracy” mode combined with either the “robust LC” or “any LC” setting. The user manual indicated that the “high accuracy” mode allows additional interference subtraction while the “robust LC” mode results in the chromatogram tails not being integrated.

We observed that the performance differences between “any LC” and “robust LC” modes were rather small (Figure 3), but there was a critical performance gap between the “high precision” and “high accuracy” modes. Furthermore, both “high accuracy” workflows were, in fact, inaccurate due to increased standard deviations (mean CV > 5 %), increased classification errors (deFDR > 1 %) and increased distortion. The “high precision” modes also resulted in higher true positive counts and performance due to the higher number of protein groups passing a 20 % CV cut-off filter.

**Figure 3:**
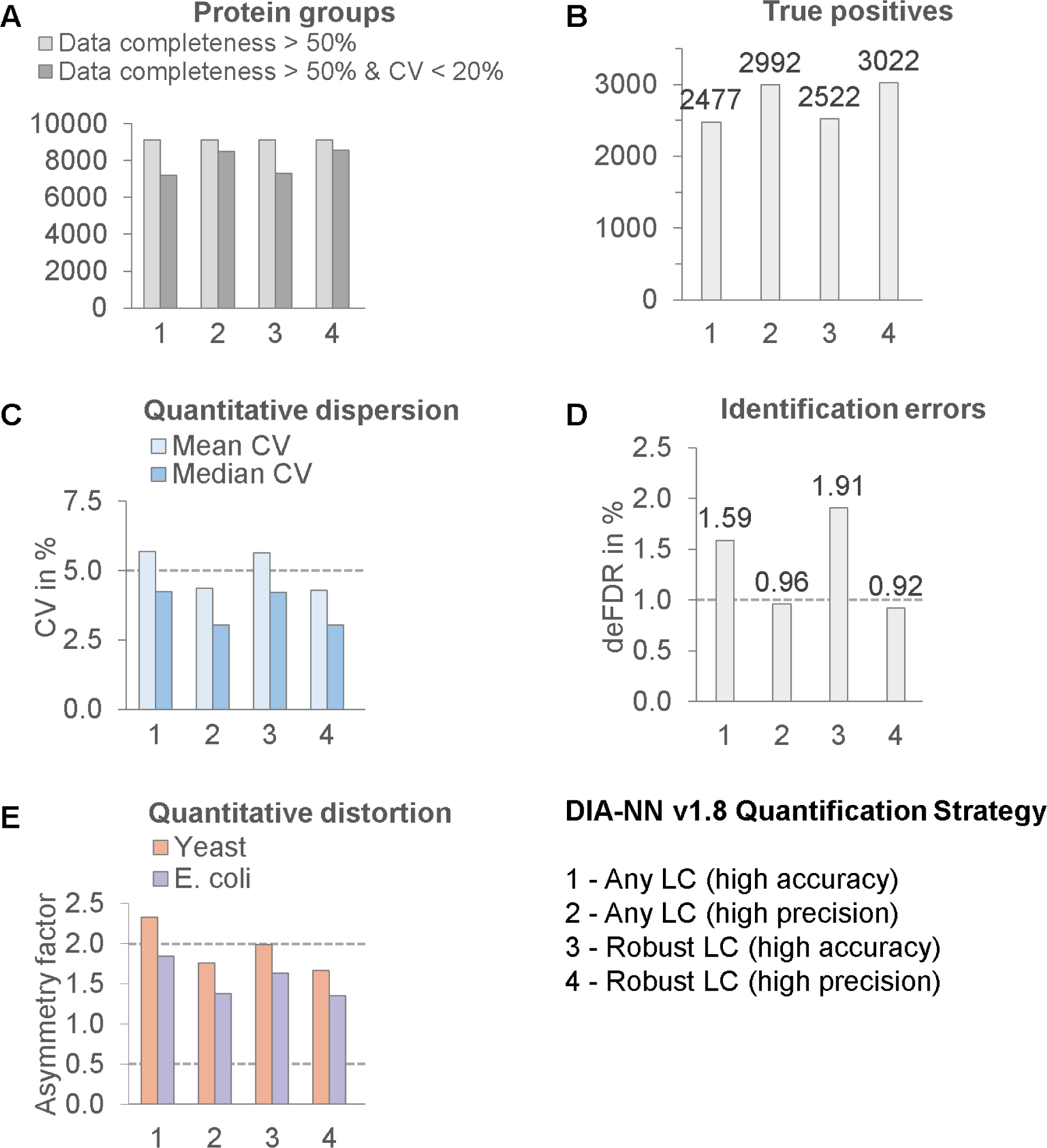
Benchmark-guided evaluation and optimization of LC-MS workflows. DIA data from QE-HF were analyzed with DIA-NN v1.8 using all 4 quantification strategies with unknown influence on quantitative accuracy (their names have to be disregarded). As shown in **C, D, E**, result sets 1 and 3 were not accurate, but 2 and 4 were. The “high accuracy” modes were found to contribute to increased classification errors, worsened precision and increased ratio expansion. Result sets 2 and 4 lead to a tolerable level of error and a higher proportion of the proteome to be precisely quantified (CV < 20 %) as shown in **A**. This translated into higher true positive values as shown in **B**. Overall, the “high precision” strategies provided improved performance and accurate quantification.

In particular, the “high precision” mode combined with the “robust LC” mode was found to be accurate and best performing, while the most commonly used default setting (“any LC - high accuracy”) reduced performance and resulted in inaccurate quantification. The optional removal of interference from the “high accuracy” modes did not provide sufficient improvement, while the drawbacks, such as reduced precision, resulted in a net negative impact on quantitative accuracy.

Our benchmark concepts were further used to determine the impact of all relevant DIA-NN settings on the QE-HF data, similar to the example above. This was crucial as the default DIA-NN settings lead to inaccurate results in all quality aspects as shown in Figure 4 C-E. A repeated stepwise optimization over two rounds was performed to maximize the number of TPs while maintaining the accuracy (Supplementary Table S2).

**Figure 4:**
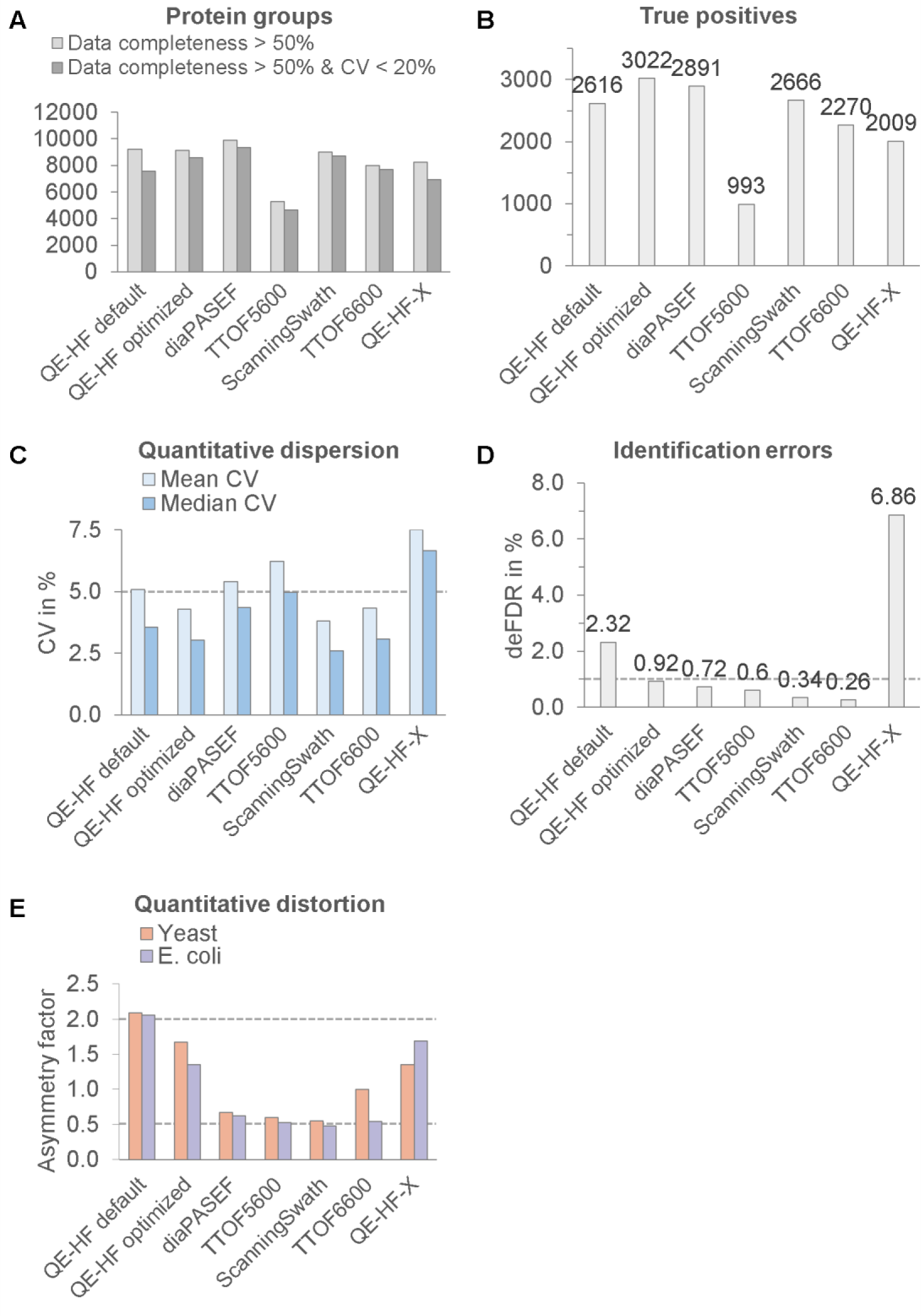
Improved cross-platform performance and accuracy assessment. QE-HF DIA data were obtained with 0.9 µg on column, 90 min gradients and both default and optimised DIA-NN v1.8 settings. Other results are from raw data of PXD028735 obtained with 1.0 or 5.0 µg on column and 120 min gradients, analysed with DIA-NN settings found to be optimal for QE-HF data. Dashed lines represent thresholds that must not be exceeded in all cases for a dataset to be considered accurate.

Changing the quantification strategy to “Robust LC - high precision”, together with the double-pass mode and a protein group identification FDR of 0.5 %, was beneficial. The corresponding results (QE-HF optimized) in Figure 4 passed all thresholds and outperformed the default DIA-NN settings in all quality and performance aspects.

### Cross-Platform Performance Evaluation

We have extended the comparison of benchmark results beyond the optimization of the DIA-NN settings to include results from a variety of instrumental setups. The respective raw data were taken from PXD028735 ^14^. The data cover the same multi-species sample types and were reanalyzed with the previously optimized DIA-NN settings. These result sets have inherent advantages over the QE-HF data due to longer gradient times (120 min vs. 90 min), higher sample loads (1.0 µg or 5.0 µg vs. 0.9 µg) and the DIA-NN double pass mode, which is beneficial but atypical for diaPASEF and similarly large raw data.

The 5 additional datasets obtained are shown in Figure 4. Only the TTOF 6600 SWATH data was found to be accurate alongside the optimized QE-HF data according to our thresholds outlined in Figure 2. Furthermore, despite the comparative advantages, no result set exceeded the performance obtained with QE-HF.

In particular, the results from QE-HF-X were found to be inaccurate, mainly due to the deFDR exceeding 6 % (Figure 4D), based on the underlying system error shown in Supplementary Figure S4. We expect this error to be difficult to detect with current conventional benchmark graphs and summary statistics outside of our benchmark procedures.

The other result sets with underlying ToF mass analyzers were found to be prone to dominant underestimation of fold changes (ratio compression) by showing asymmetry factors close to 0.5 in Figure 4E. This ratio compression is expected to be a reason for the generally low classification error rate with a deFDR below 1 % in Figure 4D.

However, due to the ratio compression and partially higher standard deviation (Figure 4 A and C) of multiple result sets such as the diaPASEF, their performance ranking according to the number of true positives would be lower than the ranking according to ID numbers (Figure 4 A and B). The optimized QE-HF data achieved the highest performance because it was less affected by the above limitations, despite the slightly lower ID numbers compared to diaPASEF.

## DISCUSSION

We have developed a benchmarking strategy and software that provides a comprehensive, practical assessment of the quantification accuracy and performance of bottom-up proteomics workflows. Our protocol is based on multi-species benchmarking analyses using confusion matrix summary statistics ^12,25^. It covers cross-run normalisation algorithms, identification errors, quantitative dispersion and distortion. In addition, unusual and unexpected errors such as system error in the QE-HF-X data and ratio expansion in the QE-HF data were successfully detected.

Our procedures revealed that for the QE-HF data the default DIA-NN settings lead to sub-optimal performance and inaccurate quantification. This is to be expected as DIA-NN was designed for short LC gradients used in conjunction with ToF MS data. QE-HF has fundamentally different characteristics in terms of chromatogram noise and interferences (Supplementary Figure S3). However, users tend to use default analysis settings and are unaware of these issues. Our benchmarking procedure provides a tool to enable critical and efficient optimization of LC-MS/MS workflows, regardless of how well any software features or unfamiliar components are documented.

The inclusion of true positive counts as a performance indicator provides a more accurate ranking of workflow performance than ID numbers by incorporating fold change over- or underestimation as well as quantitative precision. The asymmetry factors representing the distortions may be an oversimplification of the complex mathematical distributions, but we feel it is important to highlight the associated trends in a practical way with a number that subjectively matches the visual data representations. Whilst the ratio compression in the ToF data was expected, the dominant ratio expansion in the Orbitrap was unexpected and we may have uncovered a potential risk of compromised consistency across different instrument platforms.

Overall, we were pleasantly surprised by the insights and performance gains acquired from the benchmark-guided optimization. We were also able to show that even older mass spectrometers without ion mobility separation can achieve similar performance to newer mass spectrometers when using gradients of 90 minutes or longer.

The main limitation of the benchmark method used is the absence of errors related to biological variability and sample preparation. Therefore, it may be beneficial to replace the CV threshold of the accuracy criteria with an application-relevant optimization of sample preparation, chromatography and MS scan speed to maximize the number of proteins quantified with a CV below 20 %.

Further extensive use of our proposed summary statistics on newer instrument setups may be required to gain the experience to refine the proposed thresholds, especially in regards to the deFDR for ToF data. However, this work clearly demonstrated the potential of the proposed benchmark principles in workflow validation and optimization.

## CONCLUSION AND PERSPECTIVES

Benchmarks are essential tools to establish accurate and comprehensive quantification in bottom-up proteomics. However, they remain undervalued due to the lack of software and interpretation rules. While other omics disciplines are underway towards GSP standardization ^26–28^, we feel this should also become a routine in proteomics since computational methods have a decisive, but often unknown and poorly understood impact on the result quality. Our benchmark procedure allows proteomics practitioners to better understand the impact of computational algorithms, even without access to the code or understanding their mathematical background. We feel that simple and rational benchmark procedures provide impactful information on the overall quality and validity of protein quantification workflows. The proteomics community will greatly benefit from adopting these protocols.

## AUTHOR INFORMATION

### Notes

The authors declare no competing financial interest.

## ACKNOWLEDGMENTS

We are grateful to Vadim Demichev for the continuous support of DIA-NN as well as André Gohr of the MPI-CBG Scientific Computing Facility for the contributions regarding Limma. Figures were produced using the benchmark analysis script output with additions by BioRender (https://biorender.com).

## Notes

https://massive.ucsd.edu/ProteoSAFe/dataset.jsp?task=1dc758f1531148e7ba08aa7efb0519fb

### Competing Interest Statement

The authors have declared no competing interest.

## REFERENCES

(1) Gillet, L. C.; Navarro, P.; Tate, S.; Röst, H.; Selevsek, N.; Reiter, L.; Bonner, R.; Aebersold, R. Targeted Data Extraction of the MS/MS Spectra Generated by Data-Independent Acquisition: A New Concept for Consistent and Accurate Proteome Analysis. Mol. Cell. Proteomics 2012, 11 (6), 1–17.

(2) Krasny, L.; Huang, P. H. Data-Independent Acquisition Mass Spectrometry (DIA-MS) for Proteomic Applications in Oncology. Mol. Omi. 2021, 17 (1), 29–42.

(3) Pino, L. K.; Just, S. C.; MacCoss, M. J.; Searle, B. C. Acquiring and Analyzing Data Independent Acquisition Proteomics Experiments without Spectrum Libraries. Mol. Cell. Proteomics 2020, mcp.P119.001913.

(4) Zhang, F.; Ge, W.; Ruan, G.; Cai, X.; Guo, T. Data-Independent Acquisition Mass Spectrometry-Based Proteomics and Software Tools: A Glimpse in 2020. Proteomics 2020, 20 (17–18), 1900276.

(5) Kuharev, J.; Navarro, P.; Distler, U.; Jahn, O.; Tenzer, S. In-Depth Evaluation of Software Tools for Data-Independent Acquisition Based Label-Free Quantification. Proteomics 2015, 15 (18), 3140–3151.

(6) Demichev, V.; Messner, C. B.; Vernardis, S. I.; Lilley, K. S.; Ralser, M. DIA-NN: Neural Networks and Interference Correction Enable Deep Proteome Coverage in High Throughput. Nat. Methods 2020, 17 (1), 41–44.

(7) Doellinger, J.; Blumenscheit, C.; Schneider, A.; Lasch, P. Isolation Window Optimization of Data-Independent Acquisition Using Predicted Libraries for Deep and Accurate Proteome Profiling. Anal. Chem. 2020, 92 (18), 12185–12192.

(8) Derks, J.; Leduc, A.; Wallmann, G.; Huffman, R. G.; Willetts, M.; Khan, S.; Specht, H.; Ralser, M.; Demichev, V.; Slavov, N. Increasing the Throughput of Sensitive Proteomics by PlexDIA. Nat. Biotechnol. 2022.

(9) Jacome, A. S. V.; Peckner, R.; Shulman, N.; Krug, K.; DeRuff, K. C.; Officer, A.; MacLean, B.; MacCoss, M. J.; Carr, S. A.; Jaffe, J. D. Avant-Garde: An Automated Data-Driven DIA Data Curation Tool. bioRxiv 2019, 565523.

(10) Searle, B. C.; Pino, L. K.; Egertson, J. D.; Ting, Y. S.; Lawrence, R. T.; MacLean, B. X.; Villén, J.; MacCoss, M. J. Chromatogram Libraries Improve Peptide Detection and Quantification by Data Independent Acquisition Mass Spectrometry. Nat. Commun. 2018, 9 (1).

(11) Yu, F.; Haynes, S. E.; Nesvizhskii, A. I. IonQuant Enables Accurate and Sensitive Label-Free Quantification With FDR-Controlled Match-Between-Runs. Mol. Cell. Proteomics 2021, 20, 100077.

(12) Dowell, J. A.; Wright, L. J.; Armstrong, E. A.; Denu, J. M. Benchmarking Quantitative Performance in Label-Free Proteomics. ACS Omega 2021, 6 (4), 2494–2504.

(13) Navarro, P.; Kuharev, J.; Gillet, L. C.; Bernhardt, O. M.; MacLean, B.; Röst, H. L.; Tate, S. A.; Tsou, C. C.; Reiter, L.; Distler, U.; Rosenberger, G.; Perez-Riverol, Y.; Nesvizhskii, A. I.; Aebersold, R.; Tenzer, S. A Multicenter Study Benchmarks Software Tools for Label-Free Proteome Quantification. Nat. Biotechnol. 2016, 34 (11), 1130–1136.

(14) Van Puyvelde, B.; Daled, S.; Willems, S.; Gabriels, R.; Gonzalez de Peredo, A.; Chaoui, K.; Mouton-Barbosa, E.; Bouyssié, D.; Boonen, K.; Hughes, C. J.; Gethings, L. A.; Perez-Riverol, Y.; Bloomfield, N.; Tate, S.; Schiltz, O.; Martens, L.; Deforce, D.; Dhaenens, M. A Comprehensive LFQ Benchmark Dataset on Modern Day Acquisition Strategies in Proteomics. Sci. Data 2022, 9 (1), 1–12.

(15) Ammar, C.; Schessner, J. P.; Willems, S.; Michaelis, A. C.; Mann, M. Accurate Label-Free Quantification by DirectLFQ to Compare Unlimited Numbers of Proteomes. Mol. Cell. Proteomics 2023, 22 (7), 100581.

(16) Doellinger, J.; Blumenscheit, C.; Schneider, A.; Lasch, P. Optimization of Data-Independent Acquisition Using Predicted Libraries for Deep and Accurate Proteome Profiling. 2020, 1–21.

(17) Burger, B.; Vaudel, M.; Barsnes, H. On the Importance of Block Randomisation When Designing Proteomics Experiments. 2020.

(18) Amodei, D.; Egertson, J.; Maclean, B. X.; Johnson, R.; Merrihew, G. E.; Keller, A.; Marsh, D.; Vitek, O.; Mallick, P.; Maccoss, M. J. Improving Precursor Selectivity in Data-Independent Acquisition Using Overlapping Windows. J. Am. Soc. Mass Spectrom. 2019, 1–16.

(19) Adusumilli, R.; Mallick, P. Data Conversion with ProteoWizard MsConvert. In Methods in Molecular Biology; 2017; Vol. 1550, pp 339–368.

(20) Frewen, B.; MacLean, B.; Liebler, D. C.; Tomazela, D. M.; Tabb, D. L.; Finney, G. L.; Chambers, M.; MacCoss, M. J.; Shulman, N.; Kern, R. Skyline: An Open Source Document Editor for Creating and Analyzing Targeted Proteomics Experiments. Bioinformatics 2010, 26 (7), 966–968.

(21) Cox, J.; Mann, M. MaxQuant Enables High Peptide Identification Rates, Individualized p.p.b.-Range Mass Accuracies and Proteome-Wide Protein Quantification. Nat. Biotechnol. 2008, 26 (12), 1367–1372.

(22) Kong, A. T.; Leprevost, F. V.; Avtonomov, D. M.; Mellacheruvu, D.; Nesvizhskii, A. I. MSFragger: Ultrafast and Comprehensive Peptide Identification in Mass Spectrometry-Based Proteomics. Nat. Methods 2017, 14 (5), 513–520.

(23) Jupille, T.; Dolan, J.; Snyder, L.; Southern, D.; Hallenburg, K. Definition: Asymmetry factor. LC Resources, Inc. http://www.lcresources.com/resources/TSWiz/hs170.htm (accessed 2023-01-11).

(24) Fawcett, T. An Introduction to ROC Analysis. Pattern Recognit. Lett. 2006, 27 (8), 861–874.

(25) Doellinger, J.; Blumenscheit, C.; Schneider, A.; Lasch, P. Optimization of Data-Independent Acquisition Using Predicted Libraries for Deep and Accurate Proteome Profiling. 2020, 3–10.

(26) McDonald, J. G.; Ejsing, C. S.; Kopczynski, D.; Holčapek, M.; Aoki, J.; Arita, M.; Arita, M.; Baker, E. S.; Bertrand-Michel, J.; Bowden, J. A.; Brügger, B.; Ellis, S. R.; Fedorova, M.; Griffiths, W. J.; Han, X.; Hartler, J.; Hoffmann, N.; Koelmel, J. P.; Köfeler, H. C.; Mitchell, T. W.; O’Donnell, V. B.; Saigusa, D.; Schwudke, D.; Shevchenko, A.; Ulmer, C. Z.; Wenk, M. R.; Witting, M.; Wolrab, D.; Xia, Y.; Ahrends, R.; Liebisch, G.; Ekroos, K. Introducing the Lipidomics Minimal Reporting Checklist. Nat. Metab. 2022, 4 (9), 1086–1088.

(27) Burla, B.; Arita, M.; Arita, M.; Bendt, A. K.; Cazenave-Gassiot, A.; Dennis, E. A.; Ekroos, K.; Han, X.; Ikeda, K.; Liebisch, G.; Lin, M. K.; Loh, T. P.; Meikle, P. J.; Orešič, M.; Quehenberger, O.; Shevchenko, A.; Torta, F.; Wakelam, M. J. O.; Wheelock, C. E.; Wenk, M. R. MS-Based Lipidomics of Human Blood Plasma: A Community-Initiated Position Paper to Develop Accepted Guidelines. J. Lipid Res. 2018, 59 (10), 2001–2017.

(28) Bowden, J. A.; Heckert, A.; Ulmer, C. Z.; Jones, C. M.; Koelmel, J. P.; Abdullah, L.; Ahonen, L.; Alnouti, Y.; Armando, A. M.; Asara, J. M.; Bamba, T.; Barr, J. R.; Bergquist, J.; Borchers, C. H.; Brandsma, J.; Breitkopf, S. B.; Cajka, T.; Cazenave-Gassiot, A.; Checa, A.; Cinel, M. A.; Colas, R. A.; Cremers, S.; Dennis, E. A.; Evans, J. E.; Fauland, A.; Fiehn, O.; Gardner, M. S.; Garrett, T. J.; Gotlinger, K. H.; Han, J.; Huang, Y.; Neo, A. H.; Hyötyläinen, T.; Izumi, Y.; Jiang, H.; Jiang, H.; Jiang, J.; Kachman, M.; Kiyonami, R.; Klavins, K.; Klose, C.; Köfeler, H. C.; Kolmert, J.; Koal, T.; Koster, G.; Kuklenyik, Z.; Kurland, I. J.; Leadley, M.; Lin, K.; Maddipati, K. R.; McDougall, D.; Meikle, P. J.; Mellett, N. A.; Monnin, C.; Moseley, M. A.; Nandakumar, R.; Oresic, M.; Patterson, R.; Peake, D.; Pierce, J. S.; Post, M.; Postle, A. D.; Pugh, R.; Qiu, Y.; Quehenberger, O.; Ramrup, P.; Rees, J.; Rembiesa, B.; Reynaud, D.; Roth, M. R.; Sales, S.; Schuhmann, K.; Schwartzman, M. L.; Serhan, C. N.; Shevchenko, A.; Somerville, S. E.; St. John-Williams, L.; Surma, M. A.; Takeda, H.; Thakare, R.; Thompson, J. W.; Torta, F.; Triebl, A.; Trötzmüller, M.; Ubhayasekera, S. J. K.; Vuckovic, D.; Weir, J. M.; Welti, R.; Wenk, M. R.; Wheelock, C. E.; Yao, L.; Yuan, M.; Zhao, X. H.; Zhou, S. Harmonizing Lipidomics: NIST Interlaboratory Comparison Exercise for Lipidomics Using SRM 1950–Metabolites in Frozen Human Plasma. J. Lipid Res. 2017, 58 (12), 2275–2288.

