## Supporting Information for "Multi-species benchmark analysis for LC-MS/MS validation and performance evaluation in bottom-up proteomics"

### Max Planck Institute of Molecular Cell Biology and Genetics (MPI-CBG), Pfotenhauerstraße 108, 01307 Dresden, Germany

*

*

#### SUPPLEMENTARY INFORMATION

##### List of Supplementary Figures

Supplementary Figure S1: Benchmark variant to validate cross-run normalization in bottom-up proteomics by LC-MS. 2

Supplementary Figure S2: Results of the validation of normalisation algorithms in bottom-up proteomics by LC-MS. 3

Supplementary Figure S3: Manual assessment of quantitative distortions. 4

Supplementary Figure S4: Improved benchmarking procedure also allows the detection of unexpected system failures. 5

##### List of Supplementary Tables

Supplementary Table S1: Benchmark summary statistics of QE-HF data analyzed with optimized DIA-NN v1.8 settings while again cycling through DIA-NN quantification strategies. 6

Supplementary Table S2: Benchmark summary statistics of QE-HF data analyzed with default DIA-NN v1.8 settings and QE-HF and PXD0287352 data analyzed with optimized DIA-NN settings. 7

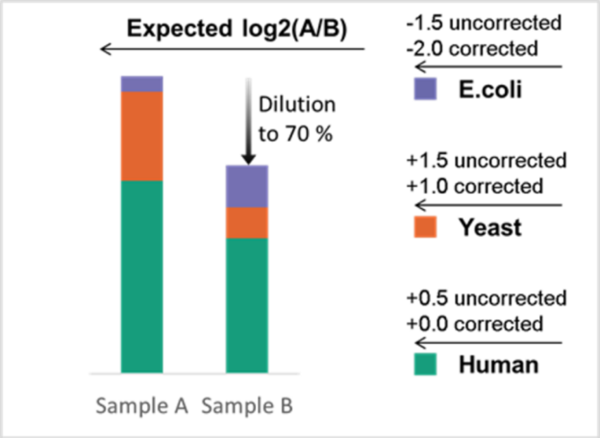


**Supplementary Figure S1: Benchmark variant to validate cross-run normalization in bottom-up proteomics by LC-MS.** Classic multi-species sample mixtures were used, but sample B was diluted to 70 % of its original concentration and load on the column. In non-normalized results, this shifts all measured log2 fold changes by approximately +0.5. Appropriate normalization would restore the originally expected log2 fold-changes of -2, 0 and +1 for *E. coli*, human and yeast, respectively. Deviations from this expectation may lead to the detection of insufficient or incorrect normalization algorithms.

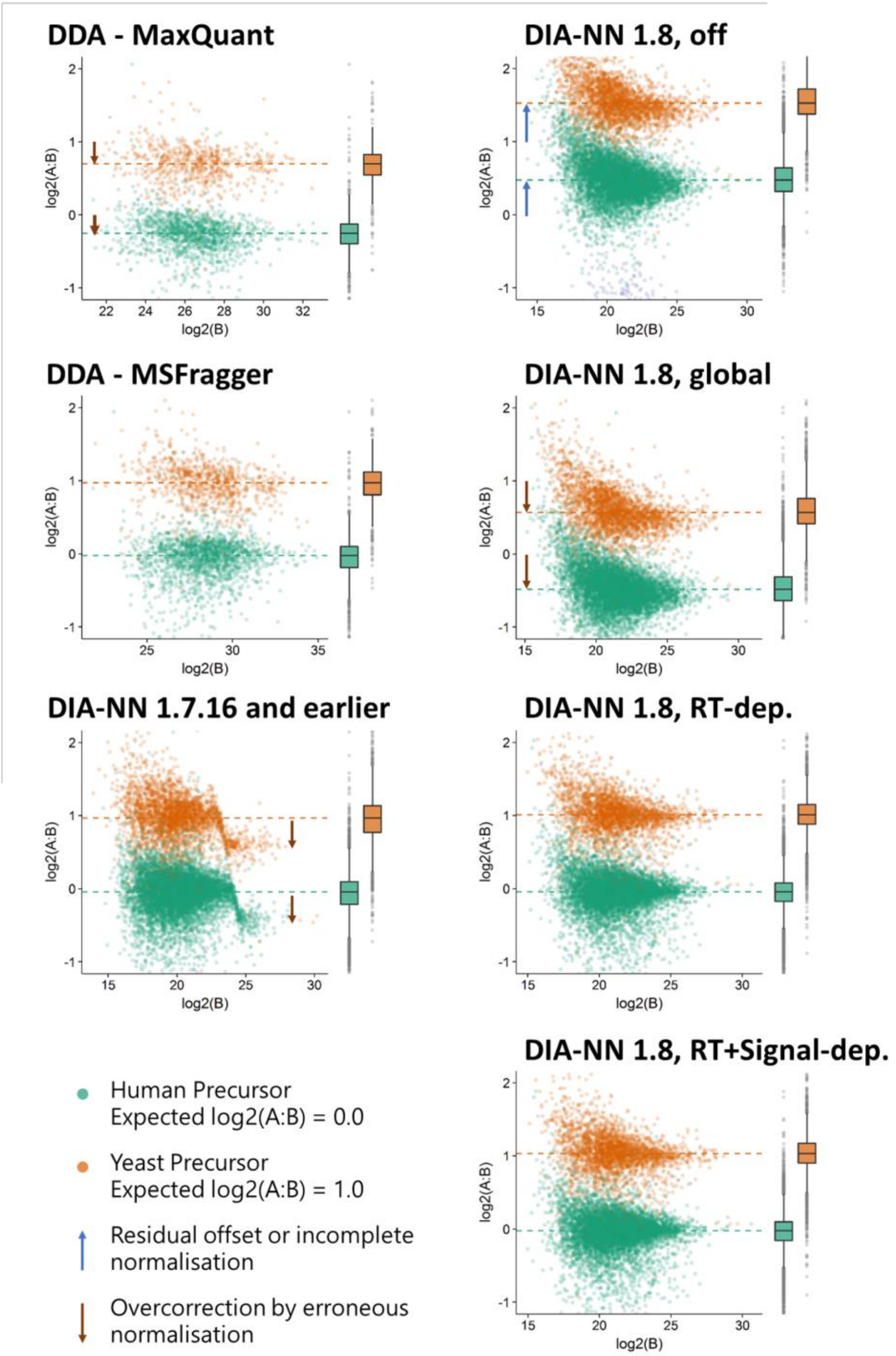


**Supplementary Figure S2: Results of the validation of normalisation algorithms in bottom-up proteomics by LC-MS.** The plots illustrate precursor quantifications with a focus on yeast and human for visualisation. Data were acquired as DIA or DDA on a QE-HF and analysed using the respective software. Dilution of benchmark sample B to 70 % results in non-normalised log2 fold changes being shifted by approximately +0.5 units to +1.5 and +0.5 for yeast and human, respectively. Appropriate normalisation would restore the expected log2 fold-changes of +1 and 0. Errors or imperfect normalisation are highlighted with the appropriate arrows. While an earlier version of DIA-NN contained a normalisation error for the most abundant precursors, the current versions of RT-dependent normalisation and MSFragger perform as desired.

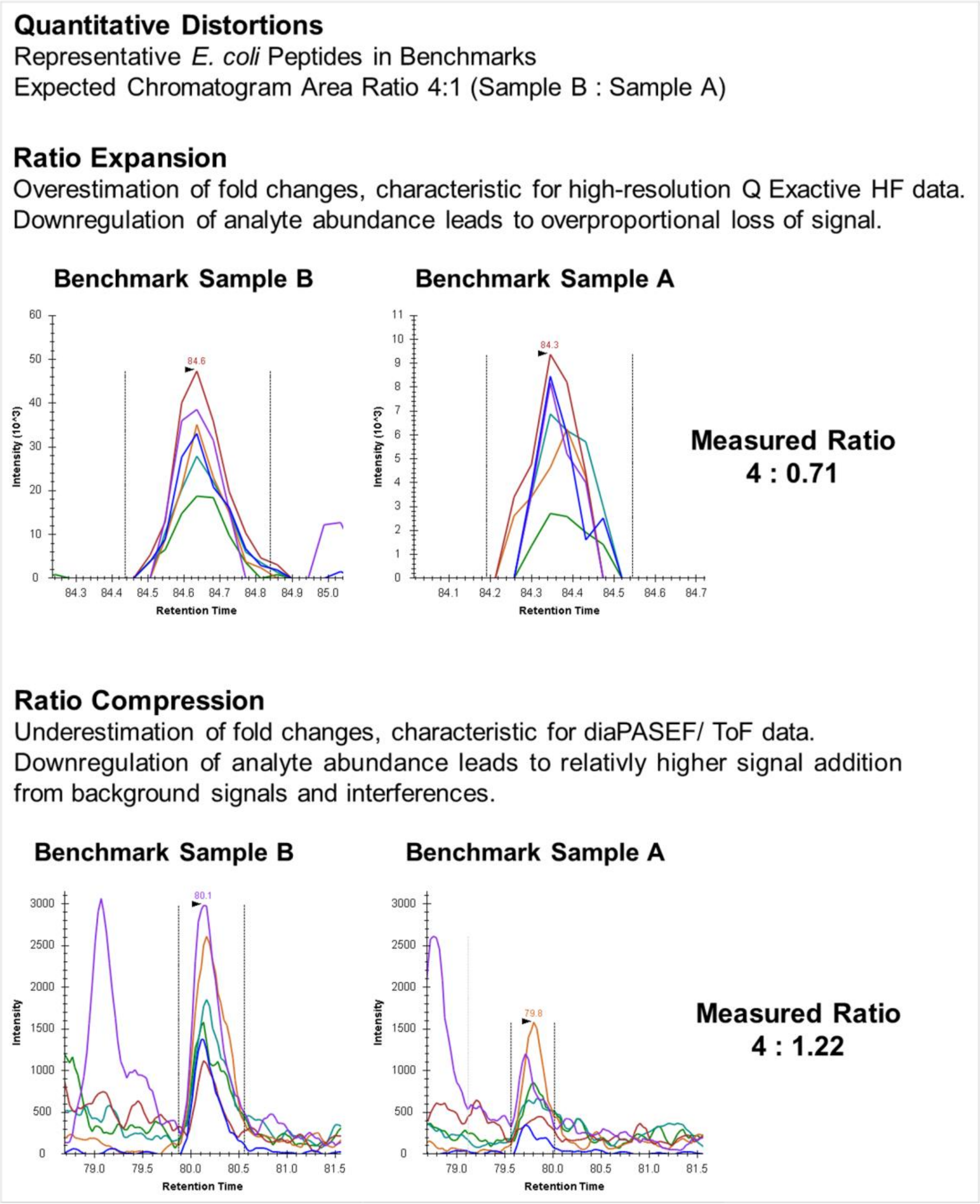


**Supplementary Figure S3: Manual assessment of quantitative distortions.** QE-HF DIA data from this study and diaPASEF data from PXD028735 were visualised in Skyline. The presence of background and interferences was clearly visible in the diaPASEF data, but was hardly observable in the QE-HF data. This is probably due to the fact that the QE-HF data are inherently subject to aggressive noise trimming and were acquired using high resolution and low-tailing chromatography (µPAC column), a 9 m/z precursor specificity for DIA after demultiplexing, and an MS2 resolution of 30000 at 200 m/z. However, low intensity signals in the QE-HF data suffered from visible loss of signal intensity and more triangular chromatogram profiles. This resulted in a disproportionate loss of signal intensity during downregulation.

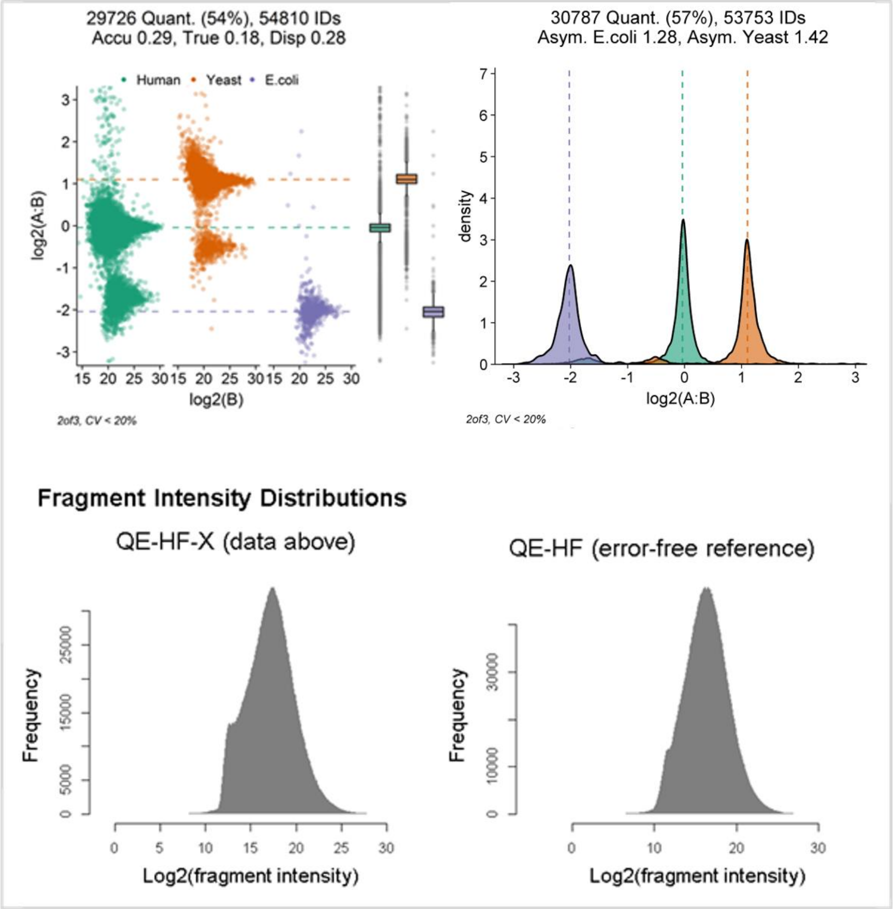


**Supplementary Figure S4: Improved benchmarking procedure also allows the detection of unexpected system failures.** The QE-HF-X data are from PXD028735 while the QE-HF data are from this study. The protein group visualisations of the QE-HF-X data are unsuspicious (not shown), but the deFDR and precursor facet plots clearly indicate the presence of a serious error. The associated offset precursor subpopulations may result from a problem possibly related to an unusually strong abnormality in the distribution of fragment intensity values.

**Supplementary Table S1:** Benchmark summary statistics of QE-HF data analyzed with optimized DIA-NN v1.8 settings while again cycling through DIA-NN quantification strategies. The "high accuracy" modes were found to produce inaccurate results, while the "high precision" modes were accurate and of higher performance.

| Result Set | Any LC (high accuracy) | Any LC (high precision) | Robust LC (high accuracy) | Robust LC (high precision) |
|---|---|---|---|---|
| deFDR | 1.59 | 0.96 | 1.91 | 0.92 |
| TP | 2477 | 2992 | 2522 | 3022 |
| Sensitivity | 99.68 | 99.77 | 99.64 | 99.74 |
| Specificity | 99.15 | 99.47 | 98.97 | 99.5 |
| Prot_ID | 9115 | 9115 | 9115 | 9115 |
| Prot_Quant | 7206 | 8505 | 7296 | 8581 |
| Prot_Asymmetry_E.coli | 2.33 | 1.76 | 1.99 | 1.67 |
| Prot_Asymmetry_Yeast | 1.84 | 1.38 | 1.63 | 1.35 |
| Prot_CV_Mean | 5.7 | 4.37 | 5.63 | 4.28 |
| Prot_CV_Median | 4.23 | 3.04 | 4.22 | 3.04 |
| Prot_Accuracy | 0.13 | 0.1 | 0.13 | 0.09 |
| Prot_Trueness | 0.33 | 0.15 | 0.3 | 0.12 |
| Prot_Dispersion | 0.1 | 0.08 | 0.11 | 0.08 |
| FP | 40 | 29 | 49 | 28 |
| TN | 4681 | 5477 | 4716 | 5523 |
| FN | 8 | 7 | 9 | 8 |
| Prec_ID | 61847 | 61854 | 61847 | 61854 |
| Prec_Quant | 37710 | 53527 | 38331 | 54397 |
| Prec_Asymmetry_E.coli | 1.63 | 1.27 | 1.65 | 1.21 |
| Prec_Asymmetry_Yeast | 1.33 | 1.27 | 1.29 | 1.07 |
| Prec_CV_Mean | 7.14 | 6.22 | 7.22 | 6.23 |
| Prec_CV_Median | 5.85 | 5.03 | 5.93 | 5.1 |
| Prec_Accuracy | 0.16 | 0.14 | 0.16 | 0.14 |
| Prec_Trueness | 0.19 | 0.11 | 0.18 | 0.09 |
| Prec_Dispersion | 0.15 | 0.13 | 0.16 | 0.14 |

**Supplementary Table S2:** Benchmark summary statistics of QE-HF data analyzed with default DIA-NN v1.8 settings and QE-HF and PXD0287352 data analyzed with optimized DIA-NN settings.

| Result Set | QE-HF default | QE-HF optimized | diaPASEF | TTOF5600 | ScanningSwath | TTOF6600 Swath | QE-HF-X |
|---|---|---|---|---|---|---|---|
| deFDR | 2.32 | 0.92 | 0.72 | 0.6 | 0.34 | 0.26 | 6.86 |
| TP | 2616 | 3022 | 2891 | 993 | 2666 | 2270 | 2009 |
| Sensitivity | 99.2 | 99.74 | 96.72 | 98.51 | 98.27 | 99.3 | 98.29 |
| Specificity | 98.74 | 99.5 | 99.67 | 99.84 | 99.85 | 99.89 | 96.96 |
| Prot_ID | 9220 | 9115 | 9893 | 5278 | 9022 | 8003 | 8225 |
| Prot_Quant | 7563 | 8581 | 9356 | 4658 | 8714 | 7692 | 6918 |
| Prot_Asymmetry_E.coli | 2.09 | 1.67 | 0.67 | 0.6 | 0.55 | 1 | 1.35 |
| Prot_Asymmetry_Yeast | 2.06 | 1.35 | 0.62 | 0.53 | 0.48 | 0.54 | 1.69 |
| Prot_CV_Mean | 5.09 | 4.28 | 5.41 | 6.24 | 3.81 | 4.33 | 7.5 |
| Prot_CV_Median | 3.56 | 3.04 | 4.36 | 4.97 | 2.59 | 3.08 | 6.67 |
| Prot_Accuracy | 0.13 | 0.09 | 0.12 | 0.09 | 0.08 | 0.07 | 0.14 |
| Prot_Trueness | 0.3 | 0.12 | 0.54 | 0.24 | 0.24 | 0.15 | 0.23 |
| Prot_Dispersion | 0.1 | 0.08 | 0.09 | 0.08 | 0.07 | 0.06 | 0.12 |
| FP | 62 | 28 | 21 | 6 | 9 | 6 | 148 |
| TN | 4864 | 5523 | 6346 | 3644 | 5992 | 5400 | 4726 |
| FN | 21 | 8 | 98 | 15 | 47 | 16 | 35 |
| Prec_ID | 63780 | 61854 | 66375 | 34846 | 77535 | 70776 | 53753 |
| Prec_Quant | 45999 | 54397 | 54005 | 27580 | 72658 | 64820 | 30787 |
| Prec_Asymmetry_E.coli | 1.93 | 1.21 | 0.74 | 0.62 | 0.7 | 0.89 | 1.28 |
| Prec_Asymmetry_Yeast | 1.44 | 1.07 | 0.55 | 0.68 | 0.58 | 1.11 | 1.42 |
| Prec_CV_Mean | 6.55 | 6.23 | 8.15 | 8.5 | 5.94 | 6.88 | 9.93 |
| Prec_CV_Median | 5.21 | 5.1 | 7.44 | 7.83 | 4.85 | 5.89 | 9.72 |
| Prec_Accuracy | 0.15 | 0.14 | 0.15 | 0.12 | 0.1 | 0.1 | 0.28 |
| Prec_Trueness | 0.17 | 0.09 | 0.47 | 0.19 | 0.21 | 0.15 | 0.16 |
| Prec_Dispersion | 0.14 | 0.14 | 0.14 | 0.12 | 0.1 | 0.1 | 0.27 |
